# BACH2 dosage establishes the hierarchy of stemness and finetunes antitumor immunity in CAR T cells

**DOI:** 10.1101/2025.08.18.670909

**Authors:** Taidou Hu, Ziang Zhu, Ying Luo, Safuwra Wizzard, Jonathan Hoar, Sejal S Shinde, Kiddist Yihunie, Chen Yao, Tuoqi Wu

## Abstract

Self-renewing stem-like T cells promote the efficacy of cancer immunotherapy and are a heterogeneous population with sub-lineages demonstrating different degrees of stemness. At the apex of this hierarchy is long-term (LT) stem-like T cells with the highest capacity of persistence, repopulation and response to immune checkpoint inhibitors (ICI). However, the pathway that establishes the hierarchy of stemness in chimeric antigen receptor (CAR) T cells and its role in antitumor efficacy of CAR T cells are unclear. Here, we demonstrate that LT stem-like differentiation and antitumor immunity of CAR T cells are dose-dependently regulated by BACH2. Pre-infusion LT stem-like CAR T cells showed epigenetic activation of BACH2 and superior antitumor response in mice and humans. After clearing tumor *in vivo*, LT stem-like T cells emerged as a CAR subset that transcriptionally and epigenetically upregulated BACH2 and downregulated TOX. Loss-of-function experiments revealed an essential role of BACH2 in the antitumor response of CAR T cells and the transcriptional program of LT stem-like CAR T cells. In exhaustion-prone GD2 CAR T cells with high tonic signaling, we showed that quantitative control of BACH2 protein level by a small molecule drug finetuned the degree of stemness and exhaustion in CAR T cells. Chemically inducing temporal BACH2 activation during manufacture of GD2 CAR T cells imprinted greater antitumor immunity against solid tumor after infusion. Together, we show that BACH2 dosage establishes the hierarchy of stem-like CAR T cells and can be temporally and tunably controlled in CAR T cells to optimize differentiation and antitumor immunity.

## Introduction

Chimeric antigen receptor (CAR) T cells have been remarkably effective in treating several types of hematological malignancies^1^. However, only a sub-population of treated cancer patients experienced durable remission^2,3^. In addition, CAR T-cell therapies have achieved limited successes in treating solid tumors^4^. Whereas antigen escape remains a major cause of relapse, increasing evidence suggests that T-cell intrinsic dysfunction, also termed exhaustion, may contribute to antigen-positive relapse after CAR T-cell therapy^5^. T cell exhaustion is induced by persistent stimulation by tumor antigen and tonic CAR signaling ^6–17^. Exhaustion, characterized by upregulation of immune checkpoints including PD1, reduced proliferation and effector function, dysregulated metabolism and impaired persistence, limits the therapeutic potential of CAR T cells^6,7,15,18,19^. Thus, to improve the efficacy of CAR T-cell therapies for cancer patients, it is critical to identify the molecular circuit that overcomes T cell exhaustion.

Exhausted T cells are now known to be a heterogeneous population containing TCF1^+^ stem-like T cells, also called progenitor exhausted T cells (Tpex) ^20–33^. In both cancer and chronic infection, stem-like T cells persist via self-renewal and replenish TCF1^-^ terminally differentiated T cells ^20–33^. Stem-like T cells exhibit superior proliferative capacity and mitochondrial fitness and are critical for the efficacy of immune checkpoint inhibitors (ICIs) ^30–36^. Signature genes of stem-like T cells are associated with favorable clinical outcomes in patients treated with CAR T cells or tumor infiltrating lymphocytes (TILs) ^14–17^. Differentiation of stem-like T cells is tightly controlled at the transcriptional and epigenetic levels by transcription factors including TCF1, TOX, BACH2, FOXP1, MYB, FOXO1, and NRF2 during cancer and infection ^20,21,24–28,33,37–45^.

Recent studies have shown that stem-like T cells are hierarchically organized with various degrees of stemness ^43,46–48^. At the apex of this system is long-term (LT) stem-like T cells. Distinct from other stem-like lineages, LT stem-like T cells are protected from the epigenetic scar of exhaustion and preserve greater potential of long-term renewal and proliferative burst in response to ICI ^43,46,47^. However, the molecular mechanism that establishes the hierarchy of stemness in T cells, especially in CAR T cells, is not completely understood.

Inducing stem-like properties in CAR T cells represents a promising strategy to improve their antitumor efficacy. Various methods have been developed to induce stem-like properties in CAR T cells including overexpressing pro-stem transcription factors or naturally occurring oncogenes ^6,41,42,49^. Yet, the T cell response requires precise temporal and quantitative control of key signaling molecules including transcription factors. The constitutive overexpression approach limits the adaptability and plasticity of T cells in response to different CAR signaling and environmental cues and may risk transformation of CAR T cells. Thus, there is a need to improve CAR T cell efficacy by achieving tunable and temporal control of the stem-like differentiation by CAR T cells. Here, we showed that BACH2 activity in pre-infusion CAR T cells was highest among LT stem-like T cells, which were associated with superior antitumor response in mice and humans. A LT stem-like CAR T cell subset developed after tumor clearance, upregulated BACH2 signature, and downregulated TOX. Loss-of-function experiments showed that BACH2 was required for the expansion of CAR T cells *in vivo* and the LT stem-like T cell program. Through tunable control of BACH2 protein level in GD2 CAR T cells with tonic signaling-driven exhaustion, we showed that the degree of stemness and exhaustion in CAR T cells is controlled by the level of BACH2. By chemically inducing BACH2 activity during manufacture of GD2 CAR T cells, we showed that temporal BACH2 activation before infusion was sufficient to imprint greater immunity against solid tumor. In summary, our study has demonstrated that BACH2 dosage establishes the hierarchy of CAR T cell stemness and can be temporally and tunably controlled to optimize the differentiation and antitumor immunity of CAR T cells.

## Results

### A LT stem-like chromatin state in pre-infusion CAR T cells exhibits epigenetic activation of BACH2

Pre-infusion CAR T cells are a heterogeneous population consisting of distinct T cell subsets that correlate with different clinical outcomes ^14–16,35^. We sought to understand how these distinct starting CAR T cell populations are programmed in transcriptional and epigenetic levels using joint single-cell chromatin accessibility and transcriptome profiles (scATAC+RNA-seq). Murine CD19-specific CAR T cells were generated from T cells of C57BL6 mice as previously described ^23,50,51^. CD8^+^ T cells sorted from pre-infusion CD19 CAR T cells were used for scATAC+RNA-seq (**Extended Data** Fig. 1a). Single-cell transcriptome and chromatin accessibility profiles of 4,860 cells were used for analysis. An average of 2,003 genes and 9,609 open chromatin regions (peaks) per cell were detected.

Unsupervised clustering partitioned pre-infusion CD8^+^ CD19 CAR T cells into four clusters based on the single-cell transcriptome (**Fig. 1a**). Cluster 0 showed a differentiated phenotype and expressed genes associated with cytotoxicity (*Gzma*, *Nkg7*), proinflammatory chemokine signaling (*Ccr2*, *Ccr5*, *Cxcr3*) and transcriptional regulators of effector differentiation (*Id2*, *Zeb2*, *Tbx21*) (**Fig. 1b,c** and **Extended Data** Fig. 1b). Top marker genes of cluster 1 included those encoding transcriptional regulators such as *Bach2*, *Foxp1*, *Lef1*, and *Foxo1* that drive differentiation of stem-like T cells ^23,41,42,52–54^ (**Fig. 1b,c** and **Extended Data** Fig. 1b). The stem-like CD8^+^ subset in pre-infusion CAR T cells also upregulated *Sell* and *Ccr7* (**Fig. 1c** and **Extended Data** Fig. 1b). In addition, two proliferating clusters were found in pre-infusion CD8^+^ CAR T cells (**Fig. 1a-c** and **Extended Data** Fig. 1c).

**Fig. 1.**
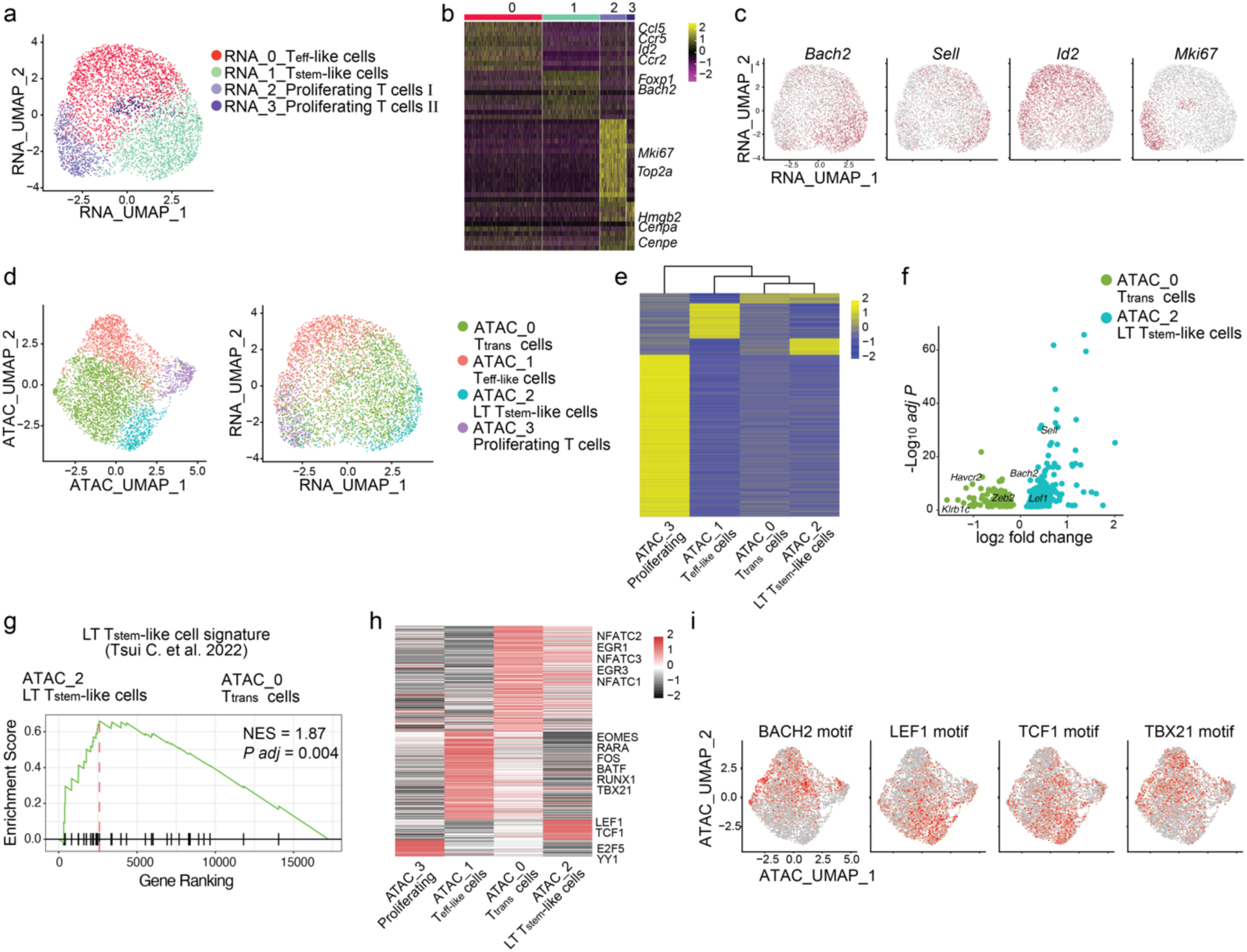
**An epigenome-defined LT stem-like subset in pre-infusion CAR T cells shows activation of BACH2. a-i**, *In vitro* expanded anti-murine CD19 CAR T cells derived from C57BL6 mice were analyzed with scATAC+RNA-seq. **a**, CD8^+^ CAR T cells are shown in a transcriptome-based UMAP plot (RNA_UMAP). Effector-like (RNA_0_Teff-like), stem-like (RNA_1_Tstem-like), and proliferating (RNA_2 and RNA_3) clusters are defined by the single-cell transcriptome and color-coded accordingly. **b**, A Heatmap of top marker genes in each transcriptome-defined cluster. Columns represent individual cells. Rows represent genes. **c**, Single-cell expression of *Bach2*, *Sell*, *Id2*, and *Mki67* shown in a UMAP plot. Color reflects the mRNA level of each gene. **d**, CD8^+^ CAR T cells projected on UMAP plots based on scATAC-seq profiles (left, ATAC_UMAP) or scRNA-seq profiles (right, RNA_UMAP). Cells are color-coded based on cluster IDs defined by the scATAC-seq profile including transitory stem-like (ATAC_0_Ttrans), effector-like (ATAC_1_Teff-like), long-term stem-like (ATAC_2_LT-Tstem-like), and proliferating (ATAC_3) clusters. **e**, A Heatmap of epigenetically activated genes in each scATAC-seq defined cluster. The color code is defined by scATAC-seq reads in the promoter and gene body. Clusters are represented by columns and organized based on hierarchical clustering. **f**, A volcano plot shows genes with differential chromatin accessibility between transitory stem-like and LT stem-like subsets. **g**, A gene-set enrichment analysis shows increased chromatin accessibility at published LT stem-like signature genes (GSE199839) in cluster ATAC_2 compared to cluster ATAC_0. NES, normalized enrichment score. *P* adj, adjusted *P* value. **h**, A heatmap of transcription factor binding motif enrichment in each scATAC-seq defined cluster. **i**, Single-cell enrichment of BACH2, LEF1, TCF1, and TBX21 (T-bet) binding motifs.

Based on unsupervised clustering of scATAC-seq profiles, pre-infusion CD8^+^ CD19 CAR T cells were segregated into four clusters and projected into either ATAC-defined uniform manifold approximation and projection (UMAP) or RNA-defined UMAP (**Fig. 1d**). scATAC-seq defined clusters showed a largely similar pattern as scRNA-seq defined cluster (**Fig. 1a,d**). Nonetheless, the stem-like CD8^+^ subset was further segregated into two subsets based on chromatin accessibility (**Fig. 1d**). One of these subsets displayed a transitory chromatin state between stem-like and effect-like subsets (**Fig. 1e,f**). The open chromatin profiles of the other stem-like subset showed epigenetic activation of the gene signature of LT stem-like T cells^43,48^ including *Bach2*, *Sell*, and *Lef1* (**Fig. 1f,g**). Compared to the transitory stem-like cells, LT stem-like CAR T cells had lower chromatin accessibility in genes associated with effector/exhaustion differentiation such as *Zeb2* and *Havcr2* (**Fig. 1e,f**). Transcription factor motif analysis revealed higher activity of T-box transcription factors such as T-bet (TBX21) in the transitory subset compared to the LT stem-like subset (**Fig. 1h**). In addition to enrichment of TCF1 and LEF1 motifs, LT stem-like CAR T cells showed higher activity of transcriptional repressor BACH2, indicated by lower chromatin accessibility at BACH2 targets (**Fig. 1h,i**). Together, a subset of pre-infusion CD8^+^ CAR T cells shows a chromatin state of LT stem-like T cells and epigenetic activation of pro-stem transcription factor BACH2.

### Pre-infusion LT stem-like CAR T cells mount greater expansion and retain stem-like phenotype in adoptive cell therapy

Expression of *Sell* was highest in the LT stem-like subset of pre-infusion CD8^+^ CAR T cells (**Extended Data** Fig. 2a). Therefore, we sought to understand the correlation of LT stem-like phenotype in pre-infusion CAR T cells with their immune response and differentiation after adoptive cell therapy. The CD62L^+^ LT stem-like subset and CD62L^-^ subset were sorted from pre-infusion CD8^+^ CD19 CAR T cells and transferred separately into mice bearing E2A-PBX1 pre-B cell acute lymphoblastic leukemia (ALL) as previously described ^23^. Seven days after infusion, we analyzed progenies of CD62L^+^ LT stem-like CAR T cells and their CD62L^-^ counterparts (**Extended Data** Fig. 2b). Notably, the expansion of LT stem-like CAR T cells after adoptive transfer was more than 2 folds higher than that of CD62L^-^ cells (**Fig. 2a**). In addition, the percentage of CD62L^+^CX3CR1^-^ stem-like subset was higher among progenies of LT stem-like CAR T cells than progenies of CD62L^-^ cells (**Fig. 2b**). Together, these results suggest that LT stem-like CAR T cells in the infusion product have greater capacity to expand and retain stemness after treating leukemia-bearing mice.

**Fig. 2.**
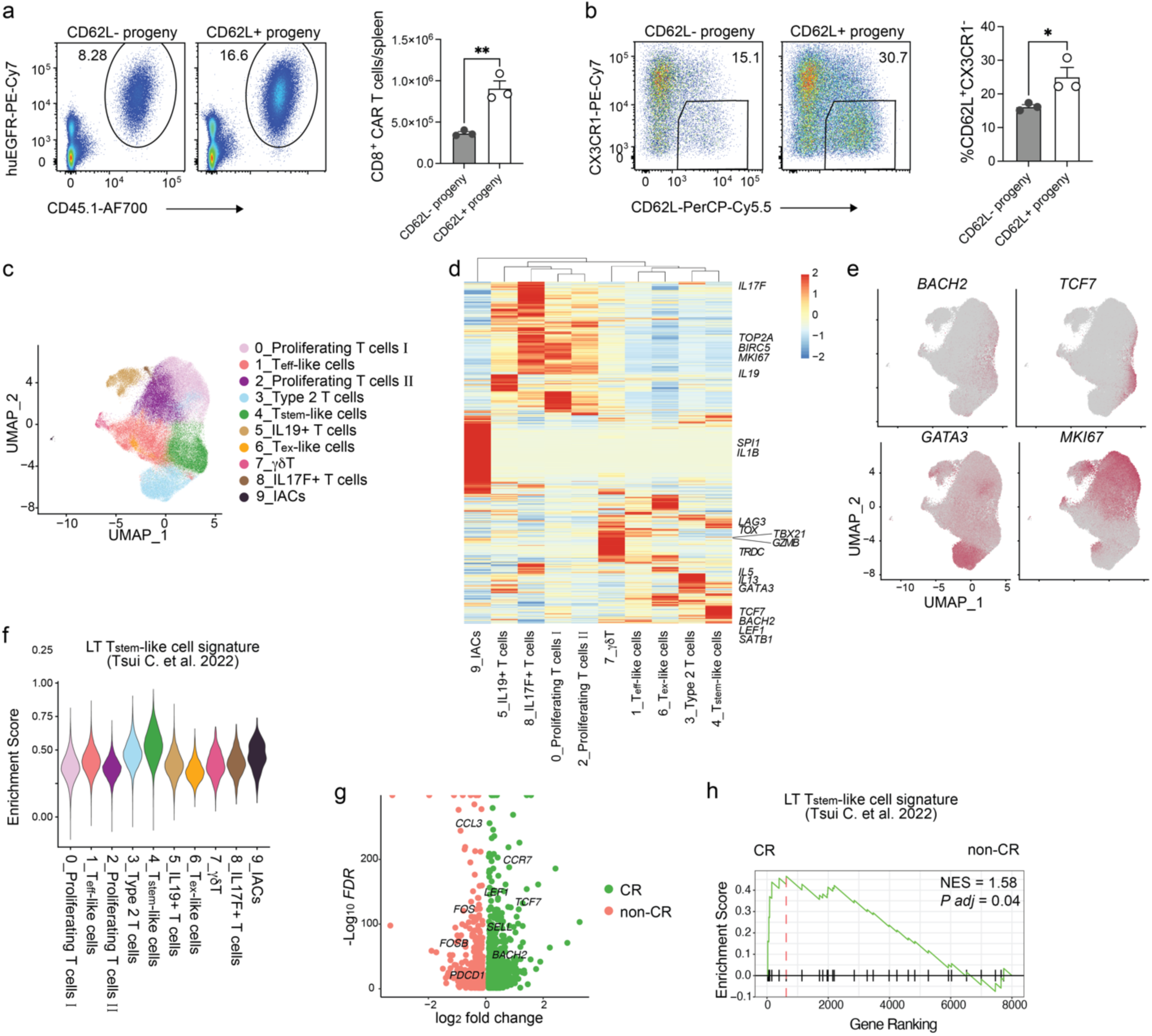
**Pre-infusion LT stem-like CAR T cells display greater antitumor responses in mice and humans. a,b**, Murine CD8^+^ CD19 CAR T cells co-expressing a truncated human EGFR (huEGFR) as a marker were sorted into CD62L^+^ and CD62L^-^ fractions, which were then transferred into C57BL6 mice bearing E2A-PBX1 B-cell ALL (n=3 mice per group). On day 7 post-transfer, representative flow cytometry plots (left) and statistics (right) of the number of CAR T cells (**a**) and percentage of CD62L^+^CX3CR1^-^ CAR T cells (**b**) derived from CD62L^+^ or CD62L^-^ pre-infusion CAR T cells are shown. **c-h**, Analysis of published scRNA-seq data (GSE241783) of CD8^+^ pre-infusion human CD19 CAR (axicabtagene ciloleucel) T cells from 40 relapsed/refractory B-cell lymphoma patients. **c**, A UMAP plot of CD8^+^ human pre-infusion CD19 CAR T cells. Cells are color-coded based on cluster IDs including proliferating T cell clusters, effector-like T cell (Teff-like) cluster, type 2 T cell cluster, stem-like T cell (Tstem-like) cluster, IL19^+^ T cell cluster, exhausted-like T cell (Tex-like) cluster, ψ8 T cell cluster, IL17F^+^ T cell cluster, and ICANS (immune effector cell-associated neurotoxicity syndrome)-associated T cell (IACs) cluster. **d**, A heatmap of differentially expressed genes in each cluster defined in panel **c**. **e**, Single-cell expression of *BACH2*, *TCF7*, *GATA3*, and *MKI67* in pre-infusion human CAR T cells. **f**, Enrichment of LT stem-like signature (GSE199839) in each cluster defined in panel **c**. **g**, A volcano plot of differentially expressed genes between CAR T cells from complete responders (CR) and those from non-CR. **h**, GSEA shows enrichment of LT stem-like signature (GSE199839) in CR versus non-CR CAR T cells. Data in **a**,**b** are representative of two independent experiments. Bar graphs represent the Mean ± Standard Error of the Mean (SEM). Circles in the bar graphs represent individual mice. Statistical significance was calculated with a two-sided Student’s t-test. \**P* < 0.05 and \*\**P* < 0.01.

### Upregulation of *BACH2* and LT stem-like signature in human pre-infusion CAR T cells of complete responders

To understand the transcriptional signature correlating with the response to CAR T cell therapy in cancer patients, we utilized single-cell RNA-sequencing (scRNA-seq) data of infusion products from 40 relapsed/refractory B-cell lymphoma patients treated with CD19 CAR T cells (axicabtagene ciloleucel (Axi-cel)) ^55^. 15 patients achieved complete response (CR), whereas 20 patients had progressive disease or partial response (non-CR). A total of 309,392 cells passed quality control with an average of 6,950 reads and 2,250 genes per cell. Unsupervised clustering identified 13 clusters including both CD8^+^ and CD4^+^ subsets (**Extended Data** Fig. 2c**-e**).

Re-clustering CD8^+^ T cells defined 10 clusters in the UMAP (**Fig. 2c**). Two proliferating clusters were identified (**Fig. 2c,d** and **Extended Data** Fig. 2f). An effector-like cluster upregulated *TBX21* and *GZMB* (**Fig. 2c,d** and **Extended Data** Fig. 2f). A type-2 CD8^+^ subset^56,57^ expressed genes associated with T_H_2 cells such as *GATA3*, *IL5*, and *IL13* (**Fig. 2c-e** and **Extended Data** Fig. 2f). We also identified a stem-like population that upregulated genes including *BACH2*, *CD27*, *TCF7*, and *LEF1* (**Fig. 2c-e** and **Extended Data** Fig. 2f). Human stem-like pre-infusion CAR T cells showed the highest level of enrichment in the LT stem-like signature compared to other subsets (**Fig. 2f**). Of note, LT stem-like signature genes including *BACH2*, *SELL*, and *TCF7* were upregulated in CD8^+^ T cells from CR patients compared to those from non-CR patients (**Fig. 2g**). In addition, gene-set enrichment analysis (GSEA) showed that LT stem-like gene signature was significantly enriched in CD8^+^ T cells from CR patients (**Fig. 2h**). Therefore, LT stem-like signature genes including *BACH2* are upregulated in human CD8^+^ T cells in the infusion product from complete responders of CD19 CAR T cell therapy.

### BACH2^high^ LT stem-like CAR T cells develop after leukemia clearance

We previously showed that treatment of CD19 CAR T cells cleared leukemia in mice bearing E2A-PBX1 B-cell ALL within seven days after infusion ^23^. To understand differentiation of CD19 CAR T cells after leukemia clearance, we performed scATAC+RNA-seq with CD19 CAR T cells from the spleen of mice inoculated with E2A-PBX1 twenty days after CAR T cell infusion. scATAC+RNA-seq profiles were generated from 4,137 CD8^+^ CAR T cells with an average of 1,928 genes and 10,941 peaks detected in each cell. Unsupervised clustering defined seven subsets based on the scRNA-seq profile (**Fig. 3a**). Among major transcriptome-defined subsets was a stem-like subset (cluster 0) that expressed high levels of *Il7r*, *Slamf6*, *Tcf7* (**Fig. 3a,b**).

**Fig. 3.**
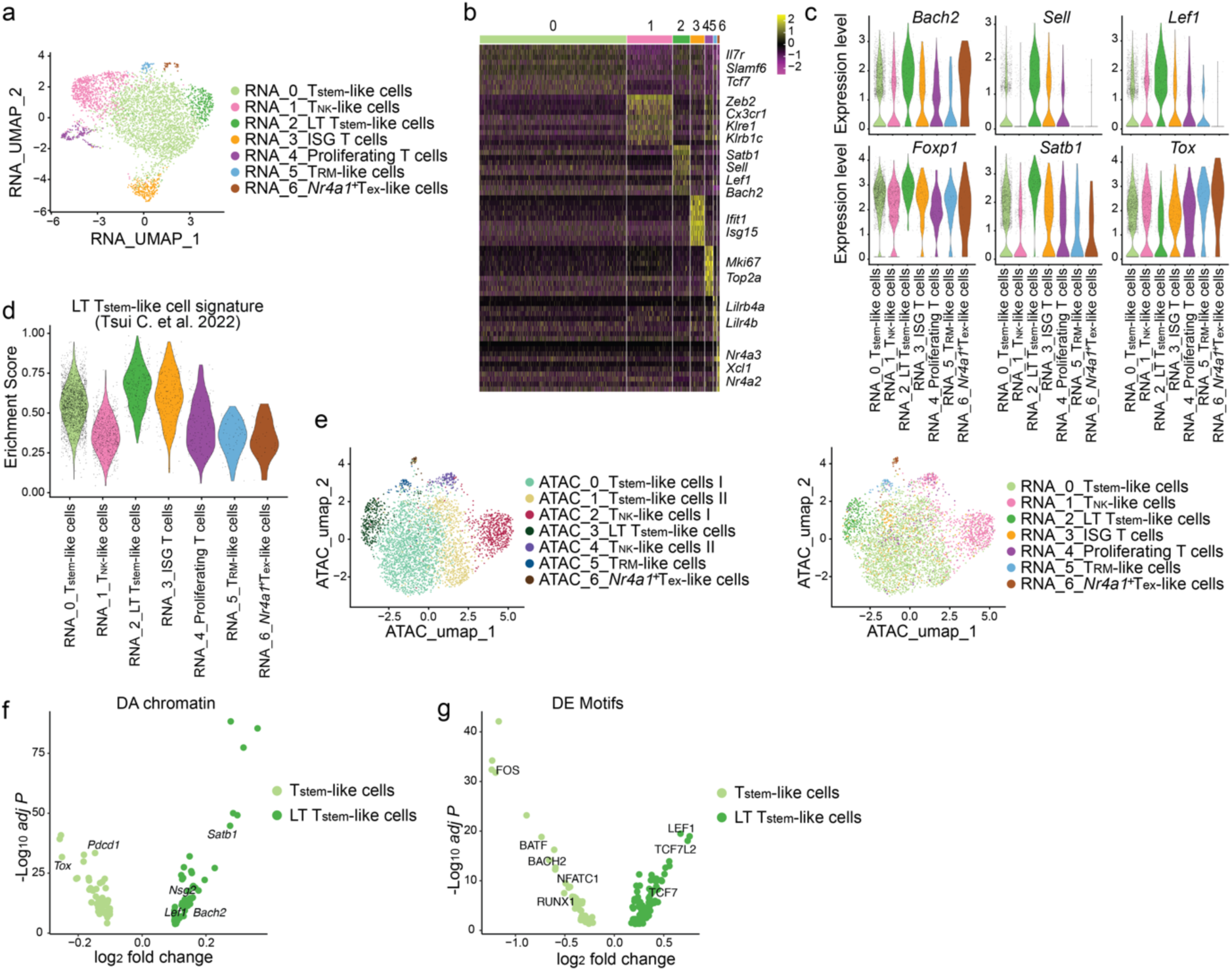
**LT stem-like CAR T cells develop after leukemia clearance and upregulate BACH2. a-g**, scATAC+RNA-Seq was performed with CAR T cells on day 20 post-treatment from C57BL6 mice that were inoculated with E2A-PBX1 and treated with CD19 CAR T cells. **a**, A transcriptome-based UMAP plot (RNA_UMAP) showing CD8^+^ CD19 CAR T cells color-coded based on cluster IDs including stem-like (RNA_0_Tstem-like), natural killer-like (RNA_1_T_NK_-like), long-term stem-like (RNA_2_LT stem-like), interferon-stimulated genes (RNA_3_ISG), proliferating (RNA_4), tissue-resident memory-like (RNA_5_T_RM_-like), *Nr4a1*^+^ exhausted-like (RNA_6_ *Nr4a1*^+^ Tex-like) clusters. **b**, A heatmap of top marker genes in each transcriptome-based cluster defined in panel **a**. **c**, Violin plots showing single-cell expression of *Bach2*, *Sell*, *Lef1*, *Foxp1*, *Satb1*, and *Tox* in each transcriptome-based cluster defined in panel **a**. **d**, Violin plots showing enrichment of LT stem-like signature in each transcriptome-based cluster defined in panel **a**. **e**, scATAC-seq based UMAP plots showing CAR T cells color-coded based on cluster IDs defined by chromatin states (left) or transcriptomes (right). **f**, A volcano plot illustrating genes with differentially accessible (DA) chromatin between stem-like and LT stem-like CAR T cells. **g**, A volcano of differentially enriched (DE) transcription factor binding motifs between stem-like and LT stem-like CAR T cells.

Cluster 1 represented a natural killer (NK)-like subset^23,58,59^ and upregulated genes including *Zeb2*, *Klre1*, *Klrb1c* (NK1.1), and *Nkg7* (**Fig. 3a,b** and **Extended Data** Fig. 3a). Notably, cluster 2 cells upregulated LT stem-like markers including *Bach2*, *Sell* (CD62L), *Foxp1*, *Lef1*, and *Satb1* and downregulated exhaustion marker *Tox* compared to stem-like CAR T cells (**Fig. 3a-c** and **Extended Data** Fig. 3b). Single-cell gene-set enrichment showed the highest enrichment of LT stem-like signature in the *Bach2*^high^ cluster 2 cells, whereas the enrichment of LT stem-like signature and the level *Bach2* transcript were intermediate in the stem-like subset and low in the NK-like subset (**Fig. 3c,d**). Thus, the transcript level of BACH2 positively correlates with the degree of stemness in CAR T cells after tumor clearance.

By connecting *cis*-regulatory elements to genes through correlating chromatin accessibility with transcription, we found that *Bach2* was among the genes controlled by the highest number of enhancers (**Extended Data** Fig. 3c). Using FigR ^60^, we connected transcription factors with their potential target genes. Notably, BACH2 was among the transcription factors with the highest number of potential targets (**Extended Data** Fig. 3d), suggesting that BACH2 is a key regulator of CAR T cells after leukemia clearance.

We next sought to understand the epigenetic program of the LT stem-like subset and stem-like subset by analyzing the scATAC-seq profile (**Fig. 3e**). The LT stem-like subset showed a chromatin state distinct from that of stem-like T cells (**Fig. 3e**). Compared to stem-like T cells, LT stem-like cells displayed greater chromatin accessibility at genes including *Bach2*, *Satb1* and *Lef1* (**Fig. 3f**). Transcription factor motif analysis revealed that compared to stem-like CAR T cells, LT stem-like CAR T cells showed higher enrichment of motifs of pro-stem transcription factors including TCF1 and LEF1 and reduced motif enrichment of AP1, NFAT, and RUNX (**Fig. 3g**). In addition, LT stem-like CAR T cells showed enhanced activity of transcriptional repressor BACH2, manifested by reduced chromatin accessibility at BACH2 binding sites (**Fig. 3g**). Therefore, our data suggest that LT stem-like CAR T cells emerge after tumor clearance and demonstrate the highest level of BACH2 activation and repression of TOX.

### After adoptive therapy a subset of CAR T cells maintains LT stem-like fate upon re-encountering tumor antigen

To evaluate the immune response and lineage stability of stem-like CAR T cells during serial encounter of tumor antigen *in vivo*, CD19 CAR T cells generated from a *Tcf7*-GFP reporter mouse^61^ were adoptively transferred into E2A-PBX1 leukemia-bearing mice. On day 7 post-infusion, *Tcf7*-GFP^+^ CD8^+^ CAR T cells and *Tcf7*-GFP^-^ CD8^+^ CAR T cells were isolated and transferred separately to a second cohort of leukemia-bearing mice. Seven days after treatment of the second cohort, ∼10-fold more progenies of *Tcf7*-GFP^+^CD8^+^ CAR T cells accumulated in the spleen and bone marrow of treated mice than progenies of *Tcf7*-GFP^-^ cells (**Fig. 4a** and **Extended Data** Fig. 4a). Whereas the majority of the progenies from *Tcf7*-GFP^+^CD8^+^ CAR T cells terminal differentiated into *Tcf7*-GFP^-^TIM3^+^, a small fraction of cells derived from *Tcf7*-GFP^+^ donors retained a *Tcf7*-GFP^+^TIM3^-^ phenotype (**Fig. 4b** and **Extended Data** Fig. 4b). In contrast, the terminally differentiated *Tcf7*-GFP^-^ donors only gave rise to *Tcf7*-GFP^-^TIM3^+^ progenies (**Fig. 4b** and **Extended Data** Fig. 4b). A small population of *Tcf7*-GFP^+^ progenies retained expression of LT stem-like marker CD62L, whereas few *Tcf7*-GFP^-^ progenies expressed CD62L (**Fig. 4c** and **Extended Data** Fig. 4c). Surface levels of PD1 and TIM3 were higher in *Tcf7*-GFP^-^ progenies than in *Tcf7*-GFP^+^ progenies (**Fig. 4d,e** and **Extended Data** Fig. 4d,e). Thus, our results demonstrate that a subset of post-treatment CAR T cells preserves proliferative potential and LT stem-like differentiation during the secondary response against tumor antigen.

**Fig. 4.**
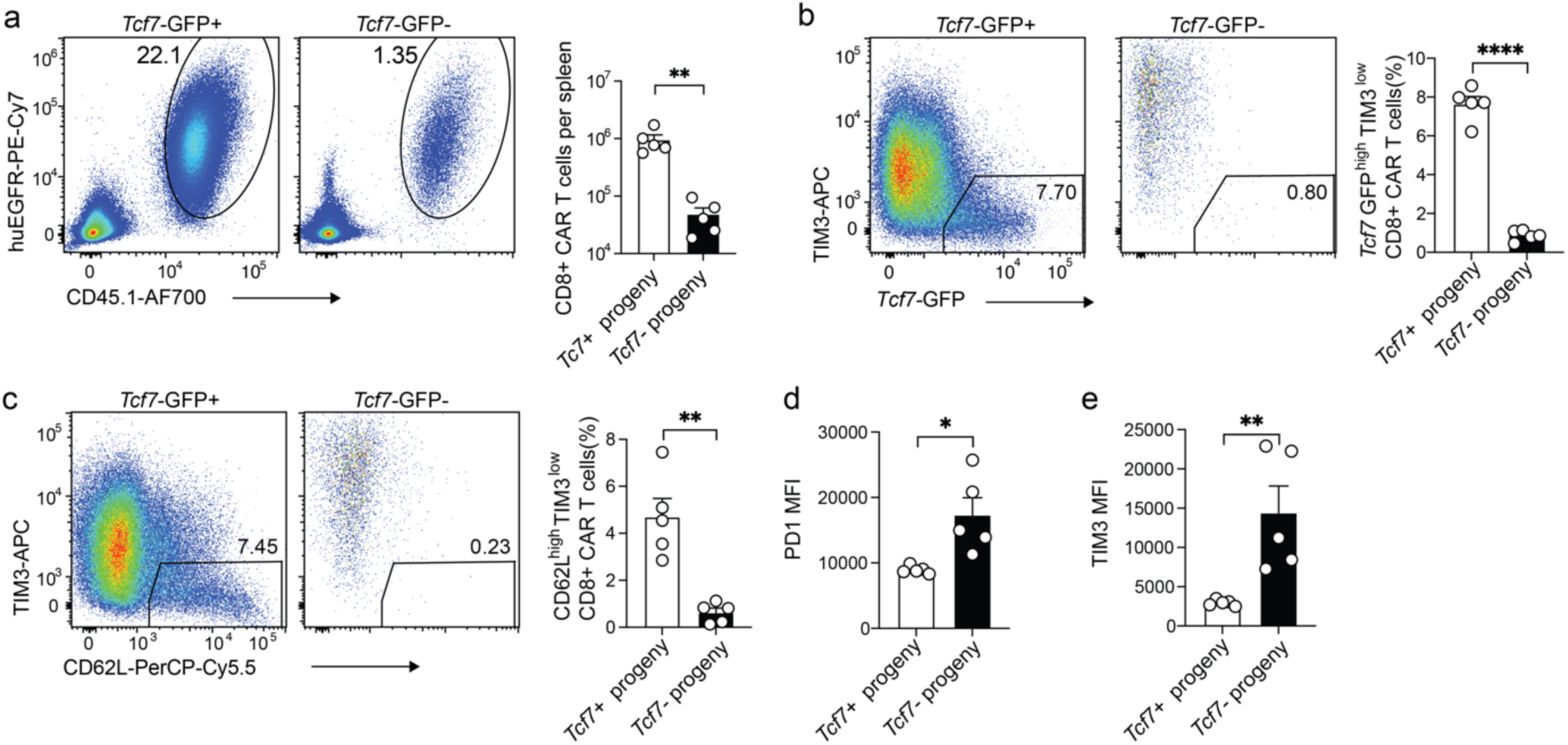
**A subset of post-infusion CAR T cells preserves LT stem-like differentiation upon tumor rechallenge. a-e**, E2A-PBX1-bearing mice were treated with CD19 CAR T cells co-expressing a truncated human EGFR (huEGFR) generated from a *Tcf7*-GFP reporter mouse. *Tcf7*-GFP^+^ and *Tcf7*-GFP^-^ CD8^+^ CAR T cells were sorted on day 7 post-infusion and transferred separately to a second batch of E2A-PBX1-bearing mice (n=5 mice per group). **a**, Left panel: Representative flow cytometry plot of splenic CD8^+^ CAR T cells (huEGFR^+^CD45.1^+^) in mice that received *Tcf7*-GFP^+^ or *Tcf7*-GFP^-^ CD8^+^ CAR T cells. Right panel: The numbers of progenies from *Tcf7*-GFP^+^ or *Tcf7*-GFP^-^ CD8^+^ CAR T cells in the spleen. **b**, Left panel: Representative flow cytometry plot of splenic *Tcf7*-GFP^high^TIM3^low^ CD8^+^ CAR T cells in mice that received *Tcf7*-GFP^+^ or *Tcf7*-GFP^-^ CD8^+^ CAR T cells. Right panel: The frequencies of splenic *Tcf7*-GFP^high^TIM3^low^ CD8^+^ CAR T cells derived from *Tcf7*-GFP^+^ or *Tcf7*-GFP^-^ donors. **c**, Left panel: Representative flow cytometry plot of splenic CD62L^high^TIM3^low^ CD8^+^ CAR T cells in mice that received *Tcf7*-GFP^+^ or *Tcf7*-GFP^-^ CD8^+^ CAR T cells. Right panel: The frequencies of splenic CD62L^high^TIM3^low^ CD8^+^ CAR T cells derived from *Tcf7*-GFP^+^ or *Tcf7*-GFP^-^ donors. **d,e**, Expression of PD1 (**d**) and TIM3 (**e**) in splenic CD8^+^ CAR T cells derived from *Tcf7*-GFP^+^ or *Tcf7*-GFP^-^ donors. Data are representative of two independent experiments. Bar graphs represent the Mean ± SEM. Circles in the bar graphs represent individual mice. Statistical significance was calculated with a two-sided Student’s t-test. \**P* < 0.05, \*\**P* < 0.01, \*\*\**P* < 0.001 and \*\*\*\**P* < 0.0001.

### BACH2 is required for the antitumor response and LT stem-like program of CAR T cells *in vivo*

To determine the role of BACH2 in the antitumor response of CAR T cells *in vivo*, we generated *Bach2*^loxP/loxP^ Cre/ERT2 CD19 CAR T cells with induced deletion of BACH2 (*Bach2* iKO) and compared them with *Bach2*^loxP/loxP^ (control) CD19 CAR T cells after adoptive transfer to E2A-PBX1-bearing mice. BACH2 deficiency led to a >2-fold reduction in the number of CD8^+^ CAR T cells and a ∼5-fold reduction in the number of CD4^+^ CAR T cells in the spleen on day 7 post-infusion (**Fig. 5a,b**). BACH2 deficiency significantly reduced the frequency of TCF1^+^ cells and the expression of LT stem-like marker CD62L in CD8^+^ CAR T cells (**Fig. 5c,d**). Immune checkpoints including TIM3 and PD1 as well as TOX were upregulated in *Bach2* iKO CD8^+^ CAR T cells (**Fig. 5e-g**). We next sought to evaluate the impact of BACH2 deficiency on the CAR T-cell response in mice bearing B16-CD19 melanoma. Similarly, BACH2 deficiency decreased the numbers of CD8^+^ CAR T cells and CD4^+^ CAR T cells in both the spleen and tumor of B16-CD19-bearing mice (**Extended Data** Fig. 5a-d). In addition, *Bach2* iKO CD8^+^ CAR T cells downregulated markers associated with stemness (**Extended Data** Fig. 5e,f). To more comprehensively understand how BACH2 regulates the differentiation of CAR T cells, we performed a scRNA-seq with 2,496 control and 4,243 *Bach2* iKO CD8^+^ CAR T cells (**Fig. 5h** and **Extended Data** Fig. 5g). BACH2 deficiency caused a clear shift in the scRNA-Seq profile of CD8^+^ CAR T cells (**Fig. 5h**), suggesting that BACH2 is a key transcriptional regulator of CD8^+^ CAR T cells. Consistent with the results above, *Bach2* iKO CD8^+^ CAR T cells contained fewer stem-like cells (cluster 1) (**Fig. 5i,j**). Comparison between control and *Bach2* iKO stem-like CD8^+^ CAR T cells showed that BACH2 deficiency downregulated stem/memory markers including *Ccr7* and *Id3* and upregulated AP1 transcription factors which are linked to exhaustion of CAR T cells ^6^ (**Extended Data** Fig. 5h). Notably, BACH2 deficient stem-like CD8^+^ CAR T cells downregulated LT stem-like gene signature compared to control stem-like CAR T cells (**Fig. 5k**). Taken together, BACH2 expression in CAR T cells is necessary for the LT stem-like transcriptional program and antitumor response.

**Fig. 5.**
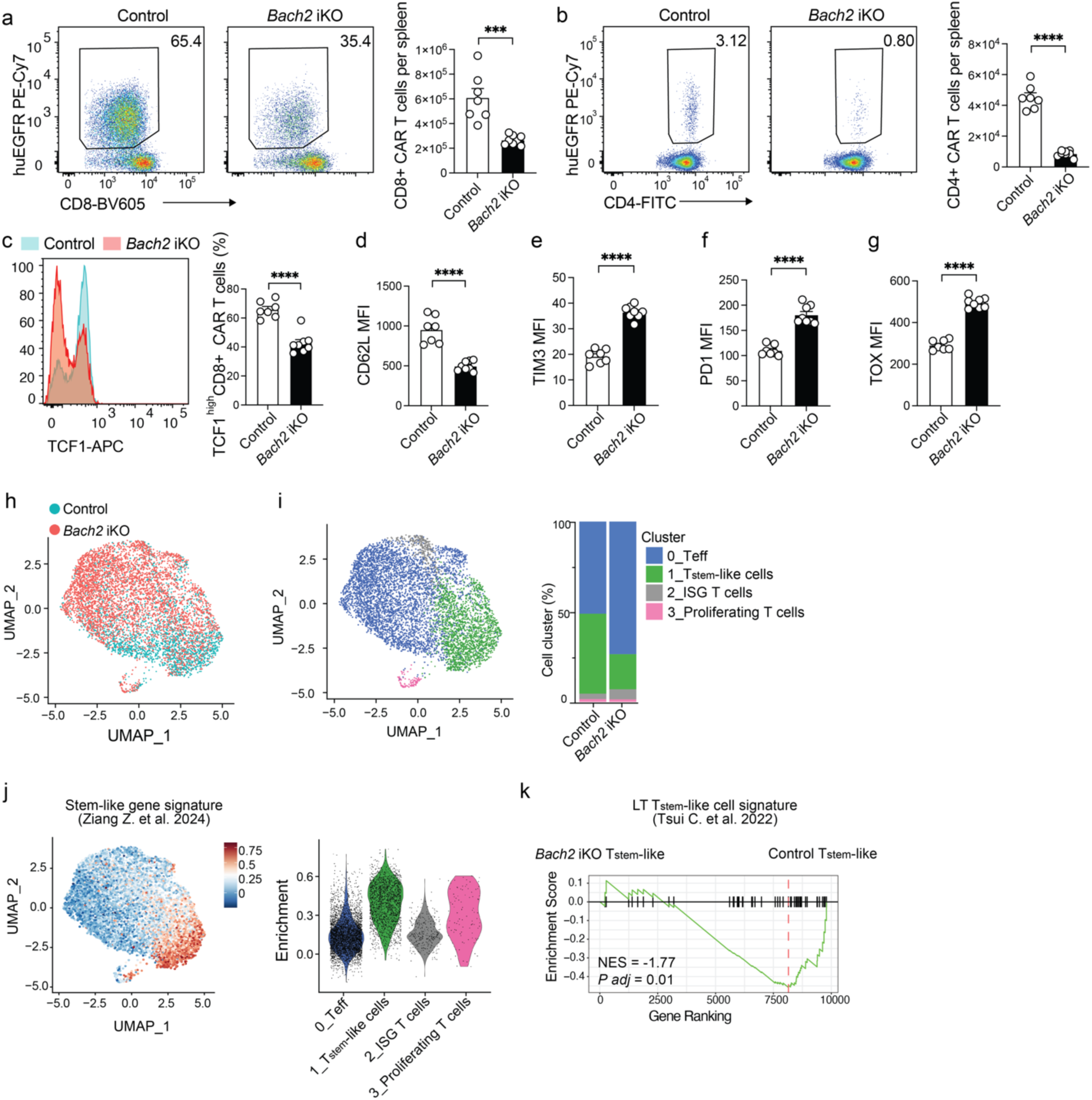
**BACH2 is a key regulator of the stemness and antitumor immunity of CAR T cells in vivo. a-g**, CD8^+^ T cells were isolated from *Bach2*^loxP/loxP^ (control) and *Bach2*^loxP/loxP^; Cre/ERT2 (*Bach2* iKO) mice after tamoxifen treatment and retrovirally transduced with CD19 CAR co-expressing a truncated huEGFR. E2A-PBX1-bearing mice treated with control or *Bach2* iKO CD19 CAR T cells were analyzed on day 7 post-infusion (control: n=7 mice, *Bach2* iKO: n=8 mice). **a**, Representative flow cytometry plots (left, gated on CD8^+^ cells) and the number (right) of splenic CD8^+^ CAR T cells (CD8^+^huEGFR^+^) in each group. **b**, Representative flow cytometry plots (left, gated on CD4^+^ cells) and the number (right) of splenic CD4^+^ CAR T cells (CD4^+^huEGFR^+^) in each group. **c**, Left panel: TCF1 expression in splenic CD8^+^ CAR T cells in each group. Right panel: The frequency of TCF1^+^ cells among splenic CD8^+^ CAR T cells in each group. **d-g**, Expression of CD62L (**d**), TIM3 (**e**), PD1 (**f**), TOX (**g**) in splenic CD8^+^ CAR T cells in each group. Data are representative of two independent experiments. Bar graphs represent the Mean ± SEM. Circles represent individual mice. Statistical significance is calculated with a two-sided Student’s t-test. \**P* < 0.05, \*\**P* < 0.01, \*\*\**P* < 0.001 and \*\*\*\**P* < 0.0001. **h-k**, B16-CD19-bearing mice were treated with control or *Bach2* iKO CD19 CAR T cells. scRNA-seq was performed with control or *Bach2* iKO splenic CD8^+^ CAR T cells on day 8 post-infusion. **h**, A UMAP plot of combined scRNA-Seq data from control or *Bach2* iKO CD8^+^ CAR T cells. Cells are color-coded based on the genotype. **i**, Left panel: A UMAP plot of control or *Bach2* iKO CD8^+^ CAR T cells color-coded based on cluster IDs. Right panel: The frequencies of cells from different clusters in control or *Bach2* iKO CD8^+^ CAR T cells. **j**, Single-cell enrichment of stem-like gene signature (GSE202543) shown in a UMAP plot (left) and violin plots (right). **k**, Enrichment of LT stem-like gene signature (GSE199839) in control stem-like (cluster 1) versus *Bach2* iKO stem-like (cluster 1) CAR T cells.

### Chemically controlling the quantity and timing of BACH2 expression in CAR T cells finetunes stem-like differentiation and optimizes tumor control

Our results above show that the quantity and timing of BACH2 expression may regulate the balance between stemness and exhaustion of CAR T cells. To finetune BACH2 expression, we fused BACH2 with a destabilizing domain (DD-BACH2). DD-fused protein is targeted for proteasomal degradation unless stabilized by a small molecule Shield-1 ^62^ (**Fig. 6a**). Varying the dose of Shield-1 tuned the amount of BACH2 protein in T cells (**Extended Data** Fig. 6a). To determine how the level of BACH2 affects the stemness and exhaustion of CAR T cells, we used an *in vitro* T cell exhaustion model in which persistent tonic signaling of GD2-specific CAR 14g2a-E101K-28z caused by antigen-independent aggregation drives T cell exhaustion ^6^. GD2 CAR T cells were co-transduced with a control construct or a construct overexpressing wildtype BACH2 or DD-BACH2. DD-BACH2 CD8^+^ GD2 CAR T cells were treated with 0nM, 100nM, or 1000nM Shield-1. Constitutive overexpressing BACH2 (BACH2 OE) increased the frequency of CD62L^+^ stem-like subset and downregulated immune checkpoints including PD1 and TIM3 (**Fig. 6b-d**). Notably, DD-BACH2 CD8^+^ GD2 CAR T cells treated with 100nM or 1000nM Shield-1 showed significantly higher frequencies of CD62L^+^ stem-like subset and lower levels of PD1 than control CD8^+^ GD2 CAR T cells (**Fig. 6b,c**). DD-BACH2 downregulated TIM3 in CAR T cells in a Shield-1 dose-dependent manner (**Fig. 6d**). Remarkably, a low level of exogenous BACH2 in untreated DD-BACH2 CD8^+^ GD2 CAR T cells was sufficient to increase the frequency of stem-like T cells and decrease PD1 and TIM3 expression albeit to a lesser extent compared to Shield-1 treated cells (**Fig. 6b-d** and **Extended Data** Fig. 6a). Thus, the degree of stemness and exhaustion in GD2 CAR T cells is quantitatively controlled by the expression level of BACH2.

**Fig. 6.**
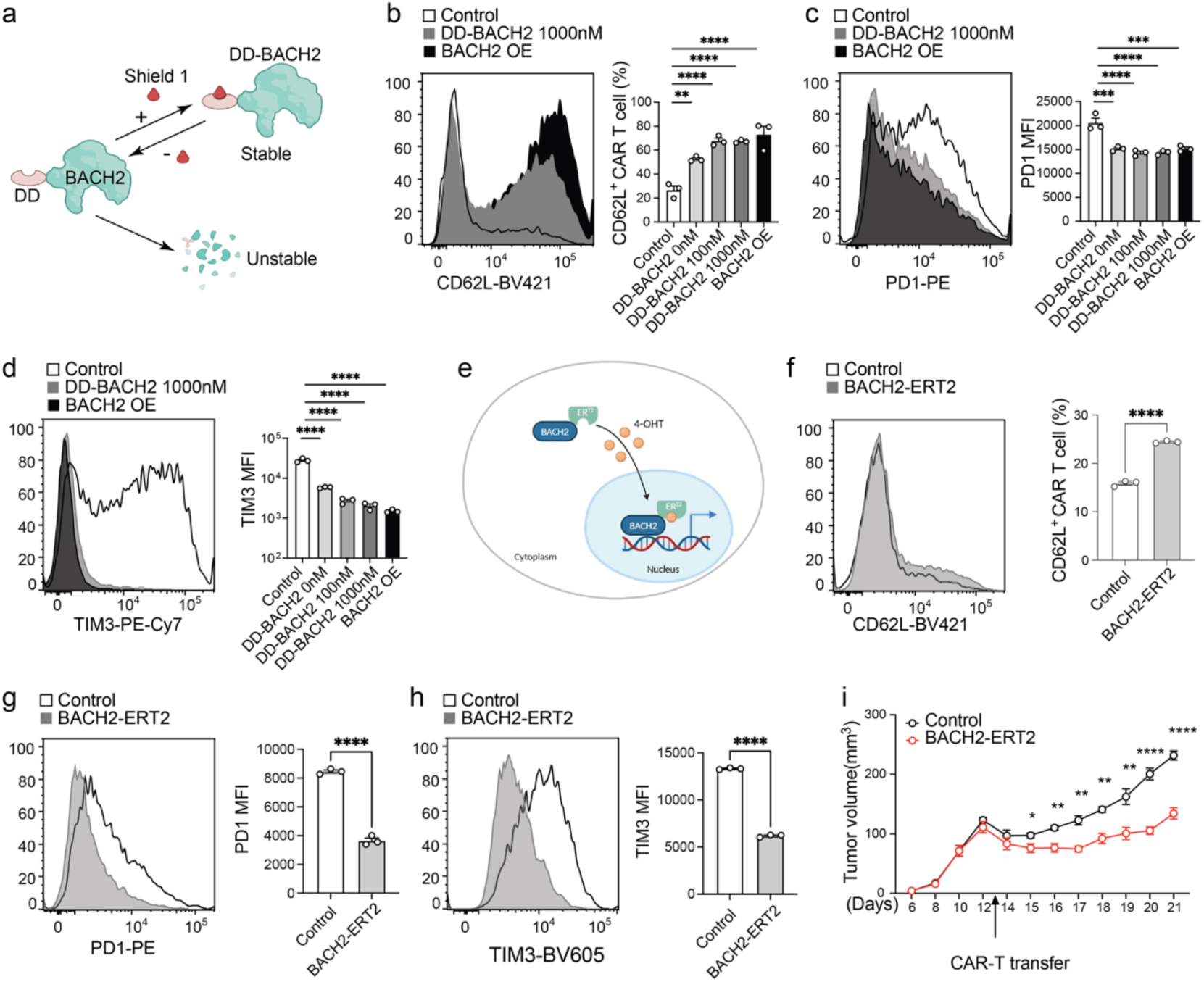
BACH2 expression quantitatively controls differentiation and antitumor immunity of CAR T cells. **a**, Schematic illustration of chemically controlling the protein level of DD-BACH2 by Shield-1. **b-d**, CD8^+^ GD2 CAR T cells were co-transduced with empty MSCV-IRES-GFP (pMIG) plasmid (control) or a pMIG plasmid overexpressing DD-BACH2 or wildtype BACH2 (BACH2 OE). DD-BACH2 GD2 CAR T cells were cultured with 0nM, 100nM, or 1000nM of Shield-1 (n=3 independent cultures per group). Expression of CD62L (**b**), PD1 (**c**), and TIM3 (**d**) in CD8^+^ CAR T cells was evaluated. **e**, Schematic illustration of inducing nuclear translocation of BACH2-ERT2 with 4-hydroxytamoxifen (4-OHT). **f-h**, CD8^+^ GD2 CAR T cells co-transduced with an empty pMIG (control) or a pMIG with BACH2-ERT2 insert were cultured with 1uM 4-OHT (n=3 independent cultures per group). The levels of CD62L (**f**), PD1 (**g**), and TIM3 (**h**) in CD8^+^ GD2 CAR T cells were determined. **i**, Tumor growth in mice bearing 9464D-GD2 neuroblastoma after treatment of control GD2 CAR T cells or BACH2-ERT2 GD2 CAR T cells described in panel **f** (n=5 mice per group). Data are representative of two independent experiments. Bar graphs represent the Mean ± SEM. Statistical significance in **b-d** was determined by one-way ANOVA with Bonferroni test to correct for multiple comparisons. Statistical significance in **f-h** was calculated with a two-sided Student’s t-test. Statistical significance in **i** was determined by two-way ANOVA with Tukey test to correct for multiple comparisons. \**P* < 0.05, \*\**P* < 0.01, \*\*\**P* < 0.001 and \*\*\*\**P* < 0.0001.

We next sought to determine whether inducing BACH2 activity during CAR T cell manufacture by controlling BACH2 nuclear translocation improves the stemness and antitumor immunity of CAR T cells. To control BACH2 nuclear translocation, BACH2 was fused to ERT2 that has enhanced reactivity to tamoxifen and minimal reactivity to endogenous estrogen as previously described ^63,64^. CD8^+^ GD2 CAR T cells co-transduced with a control construct or a BACH2-ERT2 construct were cultured with 4-hydroxytamoxifen (4-OHT) to induce nuclear translocation of BACH2-ERT2 (**Fig. 6e**). Compared to control CD8^+^ GD2 CAR T cells, BACH2-ERT2 GD2 CAR T cells showed an increase in the frequency of CD62L^+^ stem-like subset and a reduction in the levels of PD1 and TIM3 (**Fig. 6f-h**). Next, we transferred control or BACH2-ERT2 CD8^+^ GD2 CAR T cells cultured with 4-OHT into mice bearing 9464D-GD2 neuroblastoma ^45,65^ (**Fig. 6i**).

Mice treated with CD8^+^ GD2 CAR T cells programmed by BACH2 during manufacture exhibited greater tumor control than those that received control CAR T cells (**Fig. 6i**). Taken together, chemically harnessing the level and timing of BACH2 expression in CAR T cells quantitatively controls the stem-like and exhaustion differentiation programs and improves control of the solid tumor.

## Discussion

In this study, we demonstrate that the gradient of BACH2 activity establishes the hierarchy of stem-like T cells. We identified an epigenetically distinct subset of LT stem-like T cells in pre-infusion CAR T cells that showed the highest level of BACH2 activity and superior antitumor response in murine and human CAR T cells. LT stem-like CAR T cells emerged after tumor clearance and transcriptionally and epigenetically upregulated BACH2 and downregulated TOX. BACH2 deficiency in CAR T cells diminished expansion and impaired the transcriptional program of LT stem-like T cells. By chemically controlling the level of BACH2 in *in vitro* exhausted GD2 CAR T cells, we were able to calibrate the degree of stemness and exhaustion in CAR T cells. Next, we showed that GD2 CAR T cells programmed by temporally inducing BACH2 activity during manufacture exhibited greater antitumor immunity after infusion to mice with solid tumor.

Stem-like T cells were initially defined as a TCF1^+^ subset of exhausted T cells in cancer and chronic infection that self-renew, replenish TCF1^-^ terminally differentiated subsets, and respond to ICI treatment through proliferative burst^20,26–28,30–33,35^. More recently, single-cell omics revealed that stem-like T cells are a heterogenous population containing subsets with different degrees of stemness, exhaustion, and functionality. In cancer and chronic infection, a LT stem-like T cell subset has the highest capacity to persist, repopulate terminally exhausted T cells, and proliferate in response to ICI ^43,46–48^. Thus, LT stem-like T cells are at the apex of the developmental hierarchy. LT stem-like T cells also resist the exhaustion program and maintain a lower level of TOX compared to Tpex ^46^, underscoring the important clinical relevance of LT stem-like T cells. However, how the hierarchy of T cell stemness is programmed is not fully defined. Transcription factor MYB is essential for the differentiation of CD62L^+^ LT stem-like T cells, whereas MYB deficiency led to hyper-response by CD8^+^ T cells and T-cell driven immunopathology ^43^. Our results suggest that the position of T cells in the spectrum from stemness to exhaustion is determined by the level of BACH2. However, unlike MYB, loss of BACH2 did not enhance the response of terminally differentiated T cells but instead impaired expansion by CAR T cells. Therefore, BACH2 and MYB may regulate LT stem-like T cells through different downstream mechanisms. In healthy humans, two populations of stem cell-like memory T (TSCM) resemble LT stem-like T cells and Tpex, respectively ^54^. Interestingly, BACH2 is highly expressed by the activated human LT stem-like T cells, whereas activated Tpex cells express higher levels of TOX and immune checkpoints PD1 and TIGIT ^54^. Of note, our data showed that BACH2 deficiency increased TOX expression in CAR T cells. While LT stem-like T cells formed a transcriptionally and epigenetically distinct subset in CAR T cells after clearing tumor *in vivo*, the LT stem-like subset in pre-infusion CAR T cells was more distinct in the epigenetic level than the transcriptional level. Thus, it is possible that before antigen experience, a subset of pre-infusion CAR T cells is epigenetically predisposed to develop into LT stem-like T cells. Future mechanistic studies are needed to determine how the distinct chromatin states in pre-infusion CAR T cells impact their antitumor response during adoptive cell therapy.

T cell differentiation has a profound impact on their antitumor immunity ^66^. Stem-like or memory-like phenotype in premanufacture T cells or pre-infusion CAR T cells correlate with favorable outcomes in CAR T cell therapy ^14–16^. CAR T cells produced from enriched naïve/memory T cells show a greater antitumor efficacy ^35,67^. Instead of adding a cell enrichment step to the CAR T cell manufacture process, recent studies developed strategies to improve antitumor efficacy by CAR T cells through overexpressing signaling molecules that promote T cell stemness and/or repress exhaustion ^6,41,42,49^. Transcription factors dictate T-cell differentiation and function and are ideal targets for programing CAR T cells to acquire stemness and resist exhaustion.

However, instead of functioning in a binary manner, transcription factor activation is often under tight temporal and quantitative control. The precise level and timing of pro-stem transcription factors may differ for different CAR designs and tumor types. In this study, we developed two strategies to chemically control the protein level or nuclear translocation of BACH2 to allow quantitative and temporal control of CAR T cell differentiation. We further showed that temporary activation of BACH2 by a small molecule drug during manufacture of exhaustion-prone GD2 CAR T cells limited tonic signaling-driven exhaustion and enhanced control of a solid tumor. Future studies are warranted to evaluate similar strategies to control other transcriptional regulators of T cell differentiation in CAR T cells against different malignancies. In addition to improving tumor control, transcriptional regulators can be targeted in therapeutic T cells to achieve desirable cytokine profiles and/or prevent toxicity.

In summary, we have shown that the hierarchical structure of stem-like T cells is quantitatively controlled by the level of BACH2. Through controlling BACH2 activity in a temporal and tunable manner by small molecule drugs, we have engineered CAR T cells of which differentiation can be finetuned to improve tumor control. Thus, our study shed light on a new avenue to enhance the efficacy of T cell-based immunotherapy through dynamically controlling transcriptional regulators of T cell differentiation.

## Acknowledgements

The authors thank A. Guzman and A Mobley at the Flow Cytometry Core for outstanding support. We thank T. Fry (University of Colorado) for providing the MSCV-mCD19-CD28z-tEGFR plasmid and E2A-PBX1 cell line, thank C. June (University of Pennsylvania) for sharing the B16-CD19 cell line, thank C. Mackall (Stanford University) for sharing the MSGV-HA-28z (GD2 CAR) plasmid, and thank P. Sondel (University of Wisconsin) for sharing the 9464D-GD2 cell line. We thank the following funding supports: AI158294, AG083398, AG056524 from National Institutes of Health, Clinic & Laboratory Integration Program from the Cancer Research Institute, V Scholar Award from the V Foundation, and Grant for Junior Faculty from American Federation for Aging Research to T. Wu; AI154450 from National Institutes of Health, RR210035 from Cancer Prevention and Research Institute of Texas, HT94252310801 from Department of Defense to C. Yao. The funders had no influence on the design of this study, data analysis, or preparation of the manuscript.

## Contributions

Study design: T.W., C.Y. Methodology: Z.Z., T.H., Y.L., S.W., C.Y., T.W. Investigation: Z.Z., T.H., Y.L., S.W., J.H., S.S.S., K.Y., C.Y., T.W. Data curation: Z.Z., T.H., Y.L., S.W., C.Y., T.W. Writing and editing: T.W., C.Y. Visualization: T.H., C.Y. Funding acquisition: T.W., C.Y. Supervision: T.W., C.Y.

## Data availability

All data generated to support this study are available within the paper. The scRNA-Seq and scATAC+RNA-Seq data have been deposited at Gene Expression Omnibus (GSE283635). Raw and processed data for scRNA-seq of axicabtagene ciloleucel infusion products from relapsed or refractory large B-cell lymphoma patients have been published elsewhere ^55^ (GSE241783).

## Code availability

All code and software in this study are published or commercially available. Critical citations have been provided in the paper.

## Methods

### Mice

Male and female C57BL/6J (B6, strain #000664), B6.SJL-Ptprca Pepcb/BoyJ (B6 CD45.1, strain #002014), B6(Cg)-Tcf7tm1Hhx/J (*Tcf7*-GFP reporter, strain #030909) mice were purchased from the Jackson Laboratory. *Bach2*^loxP/loxP^ and *Bach2*^loxP/loxP^ Cre/ERT2 mice were described in our previous study^22^. All mice used in the experiments of this study were age and sex matched and were maintained in a B6 genetic background. Mice were kept in specific pathogen-free facilities with a 12-hour light-dark cycle. All animal procedures have been approved by the Institutional Animal Care and Use Committee at the UT Southwestern Medical Center (UTSW) under protocol numbers 103162 and 103111.

### Tumor inoculation

Syngeneic tumor cell lines E2A-PBX1 B-cell ALL, B16-CD19 melanoma, and 9464D-GD2 neuroblastoma are in a B6 background and have been described in published studies^50,65,68^. E2A-PBX1 cells were intravenously inoculated at 1×10^6^ per mouse, whereas B16-CD19 and 9464D-GD2 cells were subcutaneously injected at 1×10^6^ or 2×10^6^ per mouse, respectively.

### Transfection and retroviral transduction

For transfection of HEK293T cells, retroviral plasmid, pCL-Eco plasmid, Opti-MEM (Thermo Fisher Scientific), and TransIT-293 Transfection Reagent (Mirus Bio) were mixed, incubated and added to HEK293T cell culture. Two days after transfection, culture supernatants with retroviruses were collected and centrifuged to remove cell debris. Murine T cells activated by plate-bound anti-CD3 and anti-CD28 were transduced by a mixture of retroviruses and 8 µg/mL polybrene via spinoculation at 32°C for 90 minutes. Anti-murine CD19 CAR, described in our previous study^23^, is derived from monoclonal antibody 1D3 and contains sequences from murine CD28 and CD3ζ. An MSCV retroviral plasmid expresses anti-murine CD19 CAR and a truncated human EGFR (huEGFR) separated by a P2A peptide. An MSGV plasmid with an insert of anti-GD2 CAR was a generous gift from C. Mackall (Stanford University) and was used in our previous study ^45^. MSCV-IRES-GFP (pMIG) with a BACH2 overexpression (OE) cassette was described by us previously ^52^. A previously described BACH2-ERT2 cassette ^64^ was inserted into pMIG to generate a retroviral construct with tamoxifen responsive BACH2. To generate DD-BACH2 pMIG plasmid, a destabilization domain derived from a mutant FKBP12 ^62^ was fused to BACH2.

### Radiation and cell transfer

One day before CAR T cell transfer, mice were sub-lethally irradiated (500cGy) as previously described^23,45,51^. CD19 CAR T cells or GD2 CAR T cells were adoptively transferred into mice through tail-vein injection.

### Tamoxifen treatment

As we previously described^22^, *Bach2*^loxP/loxP^ Cre/ERT2 mice were intraperitoneally injected with tamoxifen dissolved in sunflower seed oil to induce deletion of the floxed genomic segment. Control (*Bach2*^loxP/loxP^) mice were also treated with tamoxifen.

### Flow cytometry and cell sorting

Flow cytometry and fluorescence-activated cell sorting (FACS) in this study were performed with antibodies and dyes listed below. Anti-mouse CD45.1 Alexa Fluor700 (A20, 1:200), anti-mouse CD4 FITC (RM4-5, 1:200), anti-mouse CD8a BV605 (53-6.7, 1:200), anti-human EGFR PE/Cy7 (AY13, 1:200), anti-mouse PD1 PE (RMP1-30, 1:200), anti-mouse KLRG1 BV421 (2F1, 1:200), anti-mouse TIM3 APC (RMT3-23, 1:100), anti-mouse TIM3 BV605 (RMT3-23, 1:100), anti-mouse TIM3 PE/Cy7 (RMT3-23, 1:100), anti-mouse CX3CR1 FITC (SA011F11, 1:200), and anti-mouse CX3CR1 PE/Cy7 (SA011F11, 1:800) were purchased from BioLegend. Anti-mouse CD62L PerCP/Cy5.5 (MEL-14, 1:200), anti-TOX PE (TXRX10, 1:100), Goat anti-Rabbit IgG (H+L) APC (1:500), LIVE/DEAD™ Fixable Aqua Dead Cell Staining Kit (1:250), and LIVE/DEAD™ Fixable Near-IR Dead Cell Stain Kit (1:400) were purchased from Thermo Fisher. Anti-mouse granzyme B BV421 (GB11, 1:200) was from BD Biosciences. Anti-TCF1 (C63D9, 1:200) were purchased from Cell Signaling Technology. Flow cytometry was performed with a Cytek Aurora using SpectroFlo (v3.0.3). FACS was performed with a BD FACSAria II using BD FACSDIVA (v9.0.1). Analysis was performed with FlowJo 10.10.0.

### Western blots

Proteins were extracted from the cells using RIPA Lysis and Extraction Buffer (Thermo Scientific) supplemented with Protease Inhibitor Cocktail (Thermo Scientific) and used for SDS-PAGE. The proteins were subsequently transferred to a PVDF membrane (Fisher Scientific) using a wet transfer method. The blots were blocked with 5% BSA in PBST and probed overnight at 4°C with primary antibodies against BACH2 (1:2000, Abcam, 7A4) and β-actin (1:1000, Cell Signaling Technology, 8H10D10). Following washing, the blots were incubated with secondary HRP-conjugated antibodies (Cell Signaling Technology) at a 1:5000 dilution for 60 minutes. Protein bands were visualized using enhanced chemiluminescent substrate (Thermo Scientific) according to the manufacturer’s instructions on a ChemiDoc MP Imaging System (Bio-Rad).

### Sample preparation for single-cell RNA sequencing

We generated scRNA-Seq libraries using the Chromium Next GEM Single Cell 5’ Kit (10x Genomics). Briefly, T cells labeled with TotalSeq™ Hashtag Antibody (BioLegend) from three biological replicates were sorted, which were then combined for each group. Following a wash step, cells were loaded onto a Chromium Single Cell Chip G to produce barcoded DNA. After DNA amplification, we prepared gene expression (GEX) libraries using large cDNA fragments and cell surface protein libraries (HTO) from small DNA fragments (∼200 bp). The GEX and HTO libraries were multiplexed and sequenced on an Illumina NovaSeq 6000 using the same cycle settings described above.

### Sample preparation for single-cell ATAC+RNA sequencing

We generated scRNA-Seq and scATAC-Seq libraries using the Chromium Next GEM Single Cell Multiome ATAC + Gene Expression Reagent Kits (10x Genomics), following the manufacturer’s user guide. Specifically, we sorted cells for nuclei isolation and followed the Low Cell Input Nuclei Isolation protocol from 10x Genomics to minimize nuclei loss. We tagmentated the isolated nuclei and captured them using the 10x Genomics Chromium Single Cell Controller. After GEM cleanup and pre-amplification PCR, we prepared gene expression (GEX) and ATAC libraries separately. Sequencing of the GEX libraries was performed on an Illumina NovaSeq 6000 using the following sequencing configuration: Read 1 (26 cycles), i7 Index (10 cycles), i5 Index (10 cycles), and Read 2 (90 cycles) targeting 30,000 reads per cell. The ATAC libraries were sequenced with Read 1 (50 cycles), i7 Index (8 cycles), i5 Index (24 cycles), and Read 2 (49 cycles) targeting 30,000 reads per cell.

### Data analysis for scRNA-Seq

For human CAR T cell sample, the UMI count matrix for each sample was downloaded from the GSE241783 dataset. We used R package Seurat (v4.4.0) for the subsequent analysis. We merged cells from 40 patients and filtered the cells with following criteria: 200 −7,000 detected genes, 1,000 – 15,000 detected RNA molecules and less than 15% of mitochondrial genes. To focus on CD8^+^ CAR T cells, we selected CAR T cells expressing either *CD8A* or *CD8B*, and not expressing *CD4*.

For mouse CAR T cells scRNA-Seq fastq files, we used Cell Ranger (v6.0.0) to align FASTQ files to the mm10 reference genome and to quantify barcodes and UMIs for both the gene expression and hashtag libraries with the cellranger count pipeline. We combined the cells from different samples using the merge function and retained those with 1,000–3,500 detected genes, 2,000–15,000 detected RNA molecules, and less than 5% mitochondrial gene content. After quality control, we log-normalized the data using a scale factor of 10,000. During the ScaleData function, we regressed out the effects of cell cycle, the number of detected genes, and mitochondrial gene content. TCR and IG genes were excluded from the top 2,000 variable genes identified with FindVariableFeatures, and the remaining genes were used for principal component analysis (PCA) via RunPCA. We used top 20 PCAs to calculate neighbors, clustering and UMAP. We used function FindAllMarkers or FindMarkers (min.pct = 0.1, logfc.threshold = 0.1) to determine makers genes in each cluster. We used HTODemux function to determine Hashtag antibody labeling of each biological replicate. We used pheatmap (v1.0.12) or DoHeatmap function to generate heatmaps. We calculated gene set enrichment at the single-cell level using the AddModuleScore function. Additionally, we performed gene set enrichment analysis (GSEA) for two clusters of cells using the clusterProfiler package (v4.12.6).

### Data analysis for scATAC+RNA-Seq

scATAC+RNA-Seq data were analyzed as previously described ^23^. Cell versus gene matrix of UMI counts for gene expression assay and cell versus fragment matrix for ATAC assay were generated from fastq files aligned to mm10 using cellranger-arc (v2.0.0). We performed subsequent analyses using Signac (v1) and Seurat (v4). Cells were retained if they had 1,000– 4,000 detected genes, 15,000–40,000 detected ATAC fragments, and less than 8% of UMIs mapped to the mitochondrial genome. Additionally, we excluded cells with fewer than 50% of ATAC reads mapped to peaks, a nucleosome signal greater than 2, and a TSS enrichment score below 2. Using the approach outlined in the scRNA-Seq analysis section, we analyzed the scRNA-Seq data to calculate UMAPs and identify clusters based on gene expression. For the scATAC-Seq data, peaks were called using MACS2, and those located in the mm10 genomic blacklist or nonstandard chromosomes were removed. The FeatureMatrix function was used to quantify counts within the peaks, generating a peak assay for downstream analyses. After normalizing the data, top variable features were identified using FindTopFeatures. Latent Semantic Indexing (LSI) was performed using RunSVD, and components 2 through 30 were used to calculate UMAP and identify clusters. We used geneactivity function to generate RNA activity assay and chromVAR function to generate motif assay. Differentially accessible peaks, RNA activity, and motif activity were identified using FindAllMarkers or FindMarkers. The number of correlated genes was determined using the dorcJPlot function from FigR (v0.1.0).

### Statistical analysis

For statistical analysis, Prism (GraphPad, v10.4.0) and R (v4.1.3) were used. A two-tailed Student’s t-test was used for calculating the statistical significance between two experimental conditions. A one-way or two-way analysis of variance (ANOVA) was used for determining statistical significance among more than two experimental groups. A *P* value < 0.05 was considered as statistically significant.

