## Supplemental Figures for "BACH2 dosage establishes the hierarchy of stemness and finetunes antitumor immunity in CAR T cells"

Figure S1

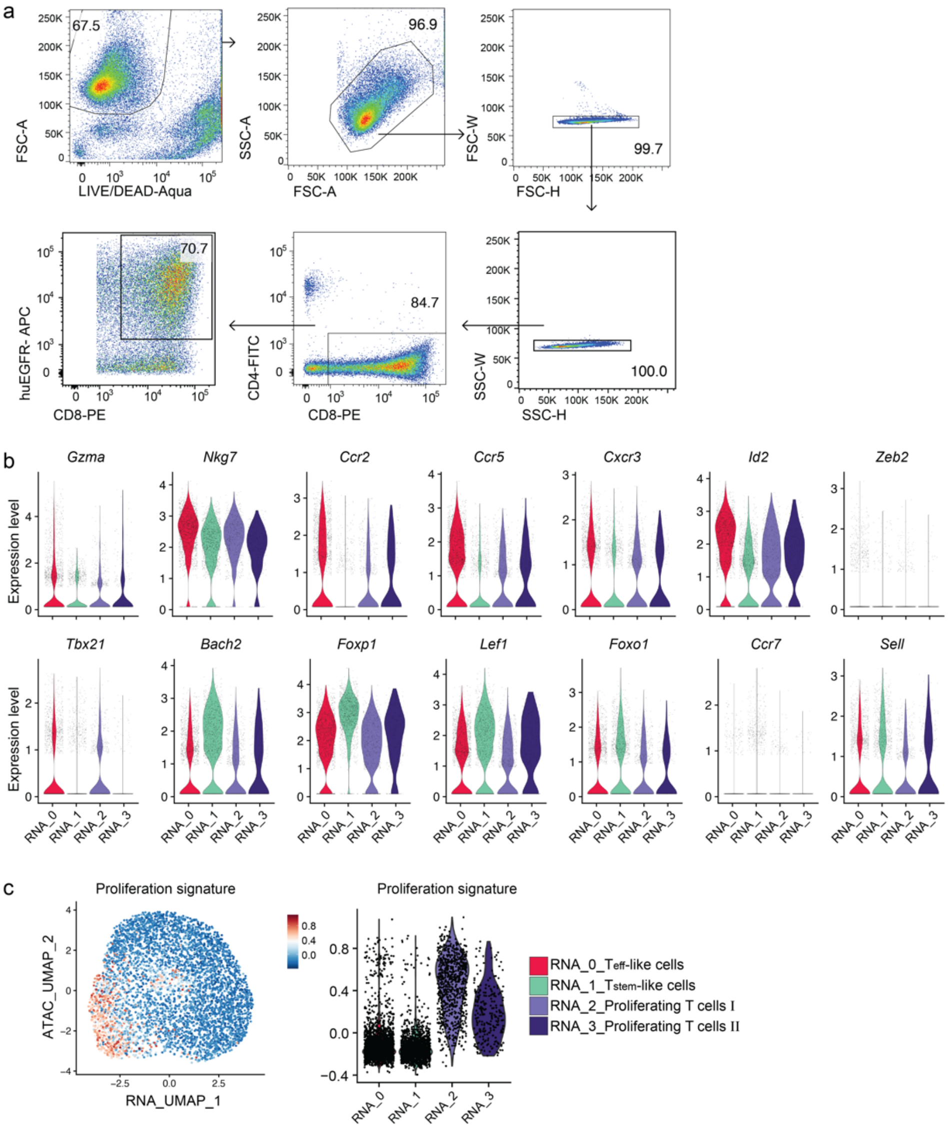


**Extended Data Fig. 1 | scATAC+RNA-seq analysis of pre-infusion anti-murine CD19 CAR T cells.**

**a,** Gating strategy for sorting *in vitro* cultured anti-murine CD19 CAR T cells. **b,c**, scATAC+RNA-seq analysis was performed as in **Fig. 1**. **b**, Violin plots showing the mRNA levels of selected genes in each transcriptome-defined cluster in **Fig. 1a**. **c**, A feature plot (left) and violin plot (right) showing the enrichment of cell-cycle gene signature in pre-infusion CD19 CAR T cells.

Figure S2

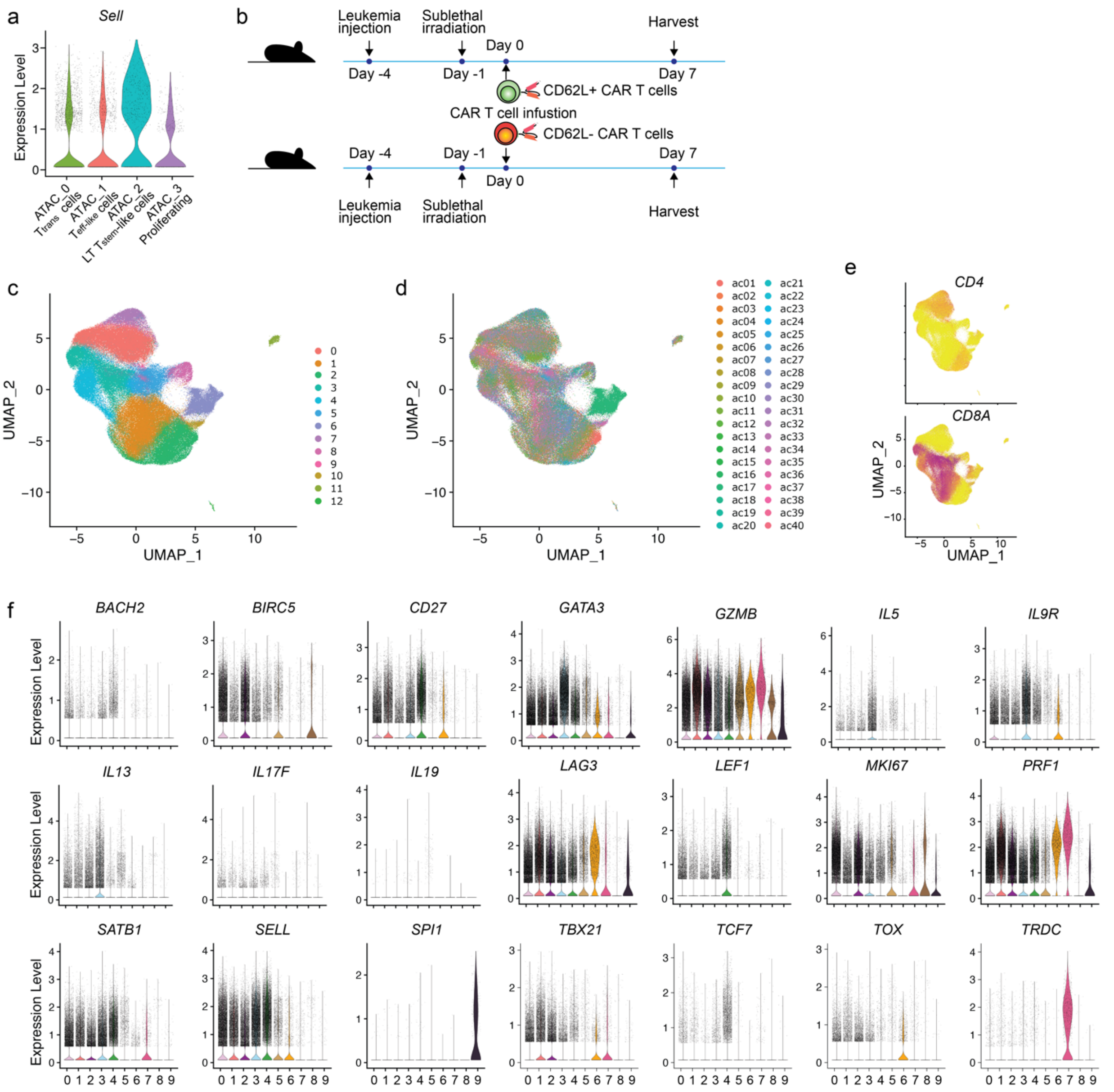


**Extended Data Fig. 2 | LT stem-like signature and *BACH2* are upregulated in pre-infusion CAR T cells from complete responders.**

**a**, Single-cell expression of *Sell* in murine pre-infusion CD8$^+$ CD19 CAR T subsets from the experiment in **Fig. 1**. **b**, Schematic illustration of the experiment in **Fig. 2a,b**. **c-f**, Re-analyzed published scRNA-seq data (GSE241783) of pre-infusion human CD19 CAR T cells from 40 relapsed/refractory B-cell lymphoma patients. **c,d**, UMAP plots of total ($CD4^+$ and $CD8^+$) pre-infusion human CAR T cells color-coded based on cluster IDs (**c**) or patient IDs (**d**). **e**, Single-cell expression of *CD4* (upper) and *CD8A* (lower). **f**, Single-cell expression of selected genes in each cluster of $CD8^+$ pre-infusion human CD19 CAR T cells. Clusters are defined in **Fig. 2c**.

#### Figure S3

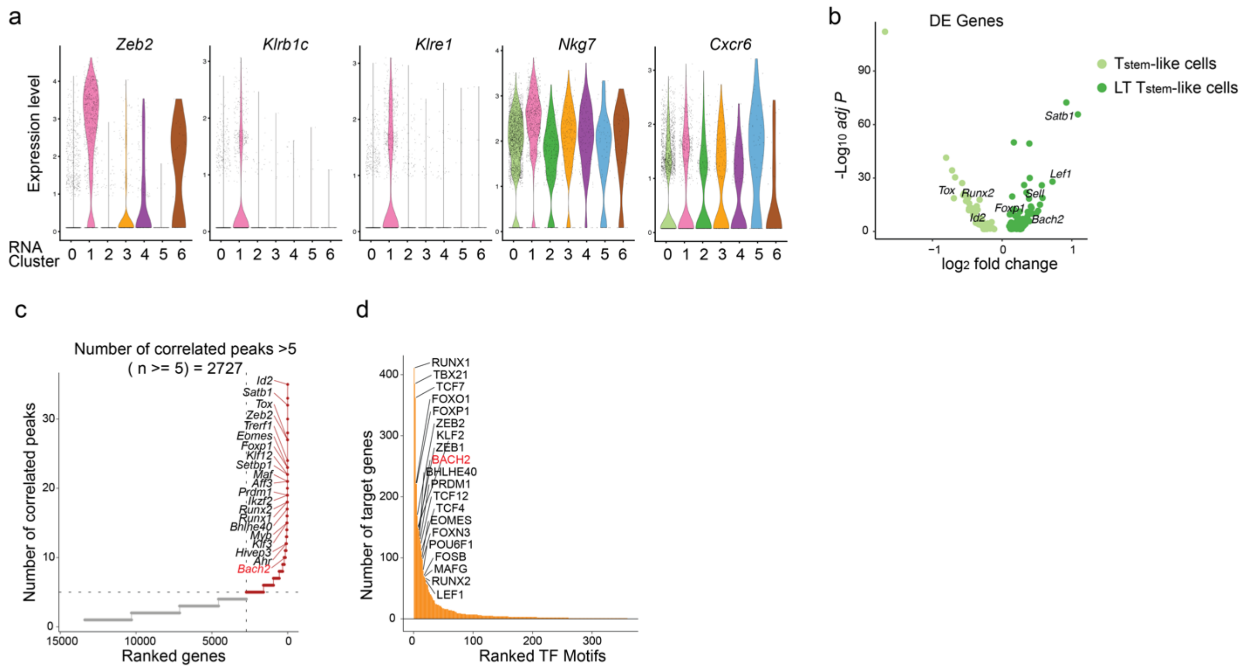


**Extended Data Fig. 3 | scATAC+RNA-seq analysis of murine CD19 CAR T cells after leukemia clearance *in vivo*.**

**a-e**, scATAC+RNA-seq experiment is described in **Fig 3**. **a**, Single-cell expression of selected genes in each transcriptome-defined cluster. **b**, A volcano plot of differentially expressed (DE) genes between stem-like and LT stem-like CAR T cells. **c**, Genes in $CD8^+$ CD19 CAR T cells are ranked by the number of correlated scATAC-seq peaks. Genes with more than five correlated peaks are highlighted in red. **d**, Transcription factors (TFs) in $CD8^+$ CD19 CAR T cells are ranked by the number of target genes.

Figure S4

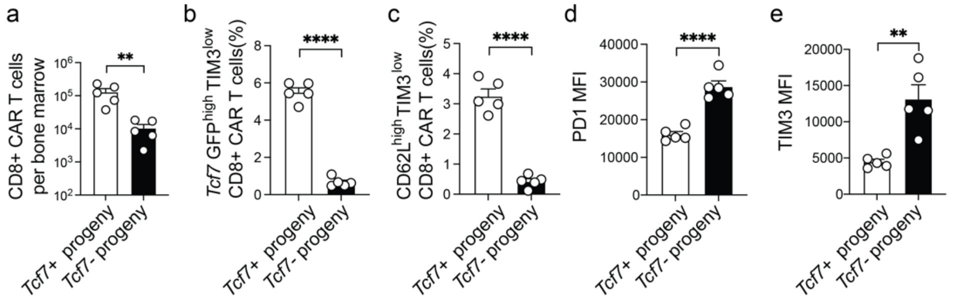


**Extended Data Fig. 4 | Antitumor response and differentiation of stem-like CAR T cells in the bone marrow during antigen rechallenge.**

**a-e**, Experimental setup is described in **Fig. 4**. **a**, The numbers of progenies from *Tcf7*-$GFP^+$ or *Tcf7*-$GFP^-$ $CD8^+$ CAR T cells in the bone marrow (n=5). **b**, The frequencies of bone marrow *Tcf7*-$GFP^{high}TIM3^{low}$ $CD8^+$ CAR T cells derived from *Tcf7*-$GFP^+$ or *Tcf7*-$GFP^-$ donors (n=5). **c**, The frequencies of bone marrow $CD62L^{high}TIM3^{low}$ $CD8^+$ CAR T cells derived from *Tcf7*-$GFP^+$ or *Tcf7*-$GFP^-$ donors (n=5). **d,e**, Expression of PD1 (**d**) and TIM3 (**e**) in bone marrow $CD8^+$ CAR T cells derived from *Tcf7*-$GFP^+$ or *Tcf7*-$GFP^-$ donors (n=5). Data are representative of two independent experiments. n, number of mice per group. Bar graphs represent the Mean ± SEM. Circles in the bar graphs represent individual mice. Statistical significance was calculated with a two-sided Student's t-test. *$P$ < 0.05, **$P$ < 0.01, ***$P$ < 0.001 and ****$P$ < 0.0001.

### Figure S5

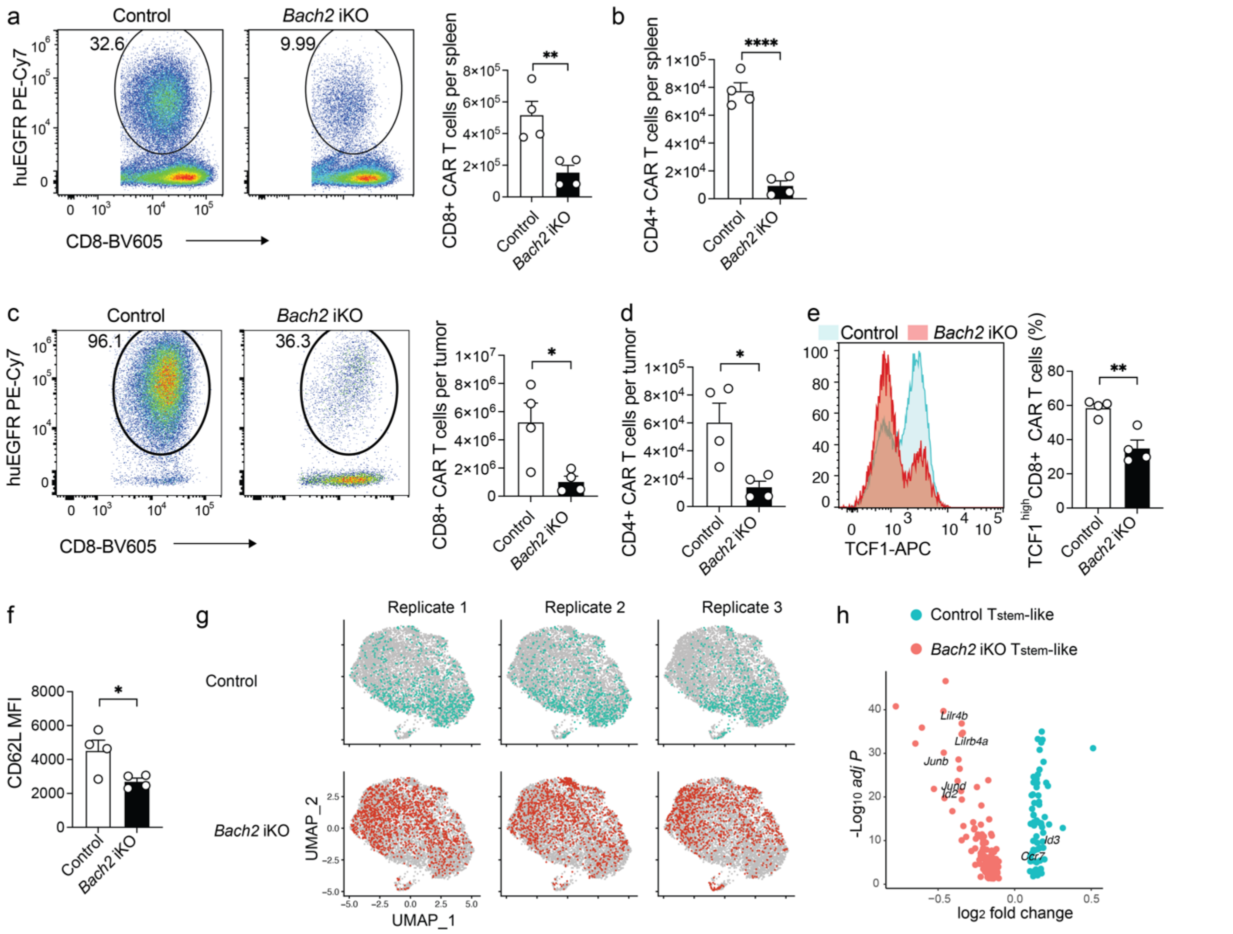


**Extended Data Fig. 5 | BACH2 deficiency impaired the antitumor response by CAR T cells.**

**a-f**, Control and *Bach2* iKO CD19 CAR T cells were generated as in **Fig. 5**. B16-CD19-bearing mice treated with control or *Bach2* iKO CD19 CAR T cells were examined on day 8 post-infusion (n=4 mice per group). **a**, Representative flow cytometry plots (left, gated on $CD8^+$ cells) and the number (right) of splenic $CD8^+$ CAR T cells ($CD8^+huEGFR^+$) in each group. **b**, The number of splenic $CD4^+$ CAR T cells ($CD4^+huEGFR^+$) in each group. **c**, Representative flow cytometry plots (left, gated on $CD8^+$ cells) and the number (right) of tumor-infiltrating $CD8^+$ CAR T cells ($CD8^+huEGFR^+$) in each group. **d**, The number of tumor-infiltrating $CD4^+$ CAR T cells

(CD4$^+$huEGFR$^+$) in each group. **e,f**, TCF1 (**e**) and CD62L (**f**) expression in control and *Bach2* iKO CD8$^+$ CAR T cells. Bar graphs represent Mean ± SEM. Circles represent individual mice. Statistical significance is calculated with a two-sided Student's t-test. *$P$ < 0.05, **$P$ < 0.01, ***$P$ < 0.001 and ****$P$ < 0.0001. **g,h**, scRNA-seq analysis of Day 8 control and *Bach2* iKO CD8$^+$ CAR T cells as in **Fig. 5**. **g**, UMAP plots of control or *Bach2* iKO CD8$^+$ CAR T cells from different individual mice (n=3 biological replicates). **h**, The volcano plot shows differentially expressed genes between control stem-like (cluster 1) and *Bach2* iKO stem-like (cluster 1) CAR T cells.

Figure S6

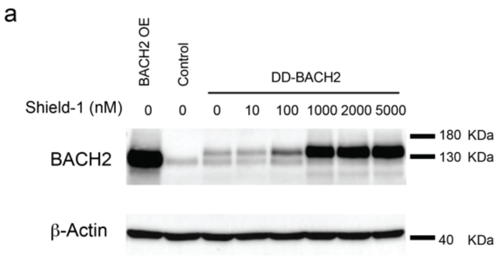


**Extended Data Fig. 6 | Control the protein level of DD-BACH2 through Shield-1.**

**a**, $CD8^+$ T cells were transduced with a pMIG plasmid (control), a pMIG-BACH2 plasmid (BACH2 OE), or a pMIG-DD-BACH2 plasmid. DD-BACH2 $CD8^+$ T cells were cultured with various concentrations of Shield-1. Western blots were used to assess the protein level of BACH2.
